# Predictable and divergent change in the multivariate P-matrix during parallel adaptation

**DOI:** 10.1101/2023.06.30.547204

**Authors:** Stephen De Lisle, Daniel I. Bolnick, Yoel E. Stuart

## Abstract

Adaptation to replicated environmental conditions can be remarkably predictable, suggesting parallel evolution may be a common feature of adaptive radiation. An open question, however, is how phenotypic variation itself evolves during repeated adaptation. Here, we use a dataset of morphological measurements from 35 populations of threespine stickleback, consisting of 16 parapatric lake-stream pairs and three marine populations, to understand how phenotypic variation has evolved during transitions from marine to freshwater environments, and during subsequent diversification across the lake-stream boundary. We find statistical support for divergent phenotypic covariance (**P**) across populations, with most diversification of **P** occurring across freshwater populations. Despite a close correspondence between within-population phenotypic variation and among population divergence, we find that variation in **P** is unrelated to total variation in population means across the set of populations. Within lake-stream pairs, however, we find that theoretical predictions for microevolutionary change can explain over 30% of the total divergence in **P** matrices across the habitat boundary. Together, our results indicate that variance evolution occurs primarily in dimensions of trait space with low phenotypic integration, driven by divergence into disparate lake and stream environments, illustrating how conserved and divergent features of multivariate variation can underlie adaptive radiation.

## INTRODUCTION

Repeated evolution of similar trait values by independent populations in similar environments provides convincing evidence for evolution by natural selection (Reznick et al. 1996, Arendt and Reznick 2006). Such parallel evolution is, therefore, an oft-invoked form of evidence for adaptation in nature (Blount et al. 2018). However, beyond the classic examples of convergence from a handful of systems for a handful of traits (*e.g.*, limb lengths in *Anolis* lizards (Losos 2009), armor loss in newly-colonizing freshwater threespine stickleback (Colosimo et al. 2005)), multivariate, multi-population convergence is more complex, and many systems show both shared and unique evolution across independent populations: a continuum of so-called (non)parallelism (Bolnick et al. 2018). Variation among populations in chance events, phenotypic plasticity, heritability, demography, environments, and in natural selection itself, generates population-specific historical contingencies (Gould 1989, Beatty 2006, 2008, Losos 2017, Blount et al. 2018). Such contingencies result in non-parallel evolutionary trajectories, even for lineages adapting to ostensibly replicated environmental gradients where parallel evolution might be expected.

Empirical evidence of (non)parallelism in trait means has become abundant (Bolnick et al. 2018, Jacobs et al. 2020, James et al. 2021, Weber et al. 2021). On the other hand, we know little about whether and why phenotypic variances and covariances may evolve predictably in these sorts of ‘natural experiments’. Yet variance-covariance structure (hereafter covariance for simplicity) has an important role in evolutionary biology. Phenotypic variation underlies the evolutionary process at all timescales, providing the material upon which natural selection acts; phenotypic variance must exist for natural selection to occur (Darwin 1859), and some of this variance must be heritable for evolutionary change to result (Fisher 1930, Lande 1976).

Thus, variance is typically viewed as a key determinant of the rate of evolution, and is often viewed as a source of constraint, biasing the direction of evolutionary change in directions of greater variability (Lande 1979, Schluter 1996, 2000, Hansen and Houle 2004). For example, under the multivariate breeders equation (Lande 1979), the evolutionary response is a product of the selection gradient **β**, and the genetic variance-covariance matrix **G** (a component of the phenotypic covariance matrix, **P**; see Glossary). If the major axis of **G** is misaligned with the direction of selection **β**, evolution should be biased away from the direction of selection and toward axes of greater covariance (Schluter 1996). However, theory predicts that natural selection can also reshape **G** and **P** (Jones et al. 2003). The speed and frequency at which this happens is the major determinant of the role of multivariate variation as constraint versus adaptation in its own right, and is an open empirical question (Svensson et al. 2021). Thus, cases where **G** or **P** is aligned with observed divergence in means may provide evidence of natural selection’s ability to shape phenotypic variation, or alternatively provide evidence of a role for variational constraints (Schluter 1996). Inferences of the shape of multivariate selection provide one step towards disentangling these interpretations (Hohenlohe and Arnold 2008, Punzalan and Rowe 2016), although estimates of contemporary selection carry their own limitations when interpreting historical adaptation (Grafen 1988). Alternatively, natural systems that show repeated adaptation to a common environmental gradient (Bolnick et al. 2018) provide an opportunity to understand the degree to which phenotypic variation itself evolves during rapid evolution, and may provide some insight into the alternative possibilities of variation as constraint vs. variation as adaptation itself (McGlothlin et al. 2018; McGlothlin et al. 2022).

Here, we use a replicated system of 16 lake-stream population pairs of threespine stickleback (*Gasterosteus aculeatus*) to quantify the predictability and repeatability of inter-population variation in the phenotypic variance-covariance matrix, **P**. Past work from this system (Stuart et al. 2017) has shown that evolutionary change across the lake-stream habitat boundary differs substantially across freshwater watersheds (i.e., the lake-stream pairs), although there is some signature of shared directions of multivariate evolution (De Lisle and Bolnick 2020). In this paper we take a comparative quantitative genetics approach to understand how multivariate phenotypic variance has changed during this radiation. Specifically, we were interested in three questions: 1) what is the extent of variation in phenotypic covariance (**P**) among habitats and populations? 2) Does within population variation captured by **P** align repeatedly with among population divergence? And, 3) can divergence in trait means predict evolution of **P** itself?

## METHODS

### Stickleback Sampling

The collection, preparation, and collation of these phenotypic data are reported in detail in (Stuart et al. 2017). Briefly, in May-July 2013, they collected adult threespine stickleback from 16 independent watersheds on Vancouver Island, British Columbia, Canada (Stuart et al. 2017). From each watershed, stickleback were sampled from one lake and its adjoining inlet or outlet stream. In addition to these 16 lake-stream pairs, they also collected marine fish from three sites spread around Vancouver Island, for a total of 35 populations (Table S1). We hereafter use the term “population” to refer to this lowest level of sampling, “pair” to refer to any individual lake-stream sample, and “watershed” to refer to the watershed from which a pair was collected. Thirty-three linear and meristic measurements, including size, were measured from each fish (Table S2). Digital landmarks were placed on left-side, lateral photographs and on ventral photographs to measure traits. Left pectoral fins were cut from each fish and splayed for photography. Standard length and a few other traits were collected via caliper or dissection. The data we use for this manuscript are a subset of those used in (Stuart et al. 2017) in that we excluded geometric morphometric data for the present study *a priori*, to ease interpretation of subsequent results.

For this study, traits were scaled through natural log transformation followed by size correction for downstream analyses. Size correction was performed by saving residuals from a linear regression of the natural log of trait value against the natural log of standard length; a single regression model for each trait was fit to maintain residual differences among populations. This trait standardization approach ensures our interpretation of changes in variance and associations with evolution of traits means (see below) is conservative, and not an inevitable outcome of mean-variance scaling. Nonetheless, similar qualitative conclusions were obtained in analyses without a log transformation.

### Trait selection

With a sample of 39 to 55 individuals per population, we lacked data to confidently estimate phenotypic covariance matrices on a population-by-population level using all 33 traits originally measured by Stuart et al. (2017), noting that a single 33- dimensional covariance matrix contains 561 unique parameters. We thus focus on a subset of traits measured, targeting seven traits because a seven-dimensional covariance matrix contains 28 unique parameters. This dimensionality ensures the possibility of robust comparison of **P** matrices among populations because the most complex model allowing among-population variation in seven-dimensional **P** still contains far fewer parameters than the number of individuals per population.

Because we were specifically interested in how variation evolves during repeatable lake-stream adaptation, we focus our analysis on the seven traits that show the most consistent change across lake-stream boundary. These seven traits were identified as the highest loading of the 32 size-corrected traits on PC1 of the among-pair correlation matrix of phenotypic change vectors describing divergence in means between lake and stream environments (Table S3 from De Lisle and Bolnick 2020); these traits were body depth, width of the pelvic girdle width, of the ventral process of the pelvic girdle (the diamond; Stuart et al. 2017), gape width, gill raker density, caudal depth, and body width. Similar qualitative conclusions were obtained using an alternative selection of seven traits based on *a priori* natural history knowledge (body depth, pelvic girdle width, gape width, gill raker number, gill raker length, dorsal spine length, pectoral fin width).

## Statistical Analysis

### Estimation of P

We used a series of multi-response mixed effects models to estimate **P** matrices and assess variation across the marine-freshwater boundary, across watersheds, and across the lake-stream boundary within pairs. In stickleback, several studies have shown that **P** and **G** align. For example, Schluter (1996) showed that the angle between the major axis of variation for a 5-trait G-matrix (**g**_max_) estimated from a limnetic freshwater population was only 16 degrees, on average, from the major axis of P-matrices (**p**_max_) calculated from several other freshwater populations. Schluter (1996) also showed that **p**_max_ and **g**_max_ made similar predictions for the observed direction of evolutionary change. Similarly, Leinonen et al. (2011) found that **p**_max_ and **g**_max_ for body shape had a correlation of r = 0.88 and an angle between them of 26 degrees, suggesting that the major axes of genetic and phenotypic variation are strongly aligned. In general, **P** and **G** are typically similar to each other (Cheverud 1988, Roff 1995, Steppan et al. 2002, Hohenlohe and Arnold 2008). However, we note that neither our analysis nor our interpretation of it depend on the substitution of **P** for **G** in that, for example, we do not use **P** to make predictions of evolutionary response.

Our mixed models were of the form

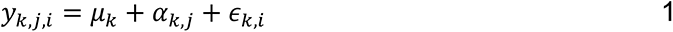

Where *y_k,j,i_* is the value of the *k*th trait from individual *i* in population *j*, *μ_k_* is the grand mean for trait *k*, *α_k,j_* is a random effect describing variation in the trait among populations, and *ϵ_k,i_* is the residual random effect describing variation among individuals. Fitting this model entails the estimation of two categories of random effect covariance matrices, the G-side covariance matrix of *α_k,j_*, which summarizes covariation in trait means across populations (i.e., the **D** matrix) (Lande 1979), and the residual covariance matrix of *ϵ_k,i_* describing among-individual variation and covariation in trait values, which is our estimate of **P**. Note that because we only have one measurement per individual fish this term will also contain measurement error, but since the fish were all measured the same way we do not expect this to contribute in a biased way to any variation in **P** we may uncover. Note also that all of our matrix comparisons accommodate uncertainty in our estimates of **P** (see below).

We fit a series of five models of the general form of equation 1 but differing in their complexity: 1) a simple model with a single **P** matrix estimated, forcing all populations and habitats to share a single within-population covariance structure, 2) a model with two separate **P**s for freshwater versus marine environments, corresponding to a shared **P**-matrix structure across all populations within each environment 3) a model with three separate **P**s for lake, stream, and marine fish, 4) a model with a separate **P** estimated for each watershed (19 **P**s; each marine population treated as its own watershed), and finally 5) a ‘saturated’ model with a separate **P** estimated for each population (habitat type x watershed combination and the three marine populations, 35 **P**s).

Finding support for the saturated model, and because we were specifically interested in potential replicated divergence in **P** matrices between lake and stream habitats, we then fit separate linear mixed models for each freshwater pair, of the form

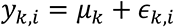

where we compared a model with separate residual covariance matrices of *ϵ_k,i_* for lake and stream environments to a reduced model with a common within-pair **P** matrix. This allowed us to assess statistical significance of lake-stream divergence in **P** for each freshwater watershed. All models were fitted by MCMC using uninformative priors in MCMCglmm (Hadfield 2010), and Deviance Information Criteria (DIC) were used to rank candidate models. Model convergence was confirmed by lack of trends in the Markov chain, as well as low estimated autocorrelation in the posterior, for both the simple and saturated models. All of the subsequent matrix comparisons (described below) were performed on the posterior distributions of **P** to account for uncertainty in our estimates.

### Describing variation in P

While the model comparison approach above can indicate whether there is statistical support for variation in **P**, other multivariate approaches are required to understand the nature of any variation that is found. We took two general approaches.

First, we performed pairwise comparisons between matrices of interest by calculating (i) the vector correlation of the leading eigenvector of two matrices, and (ii) Krzanowski’s shared subspace (Krzanowski 1979), which identifies the degree to which the parts of multivariate trait space that contain most of the variation are shared between two matrices (Aguirre et al. 2014). This is calculated by summing the eigenvalues of **S**, where

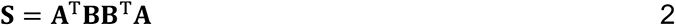

and where **A** and **B** are matrices that contain a subset of eigenvectors of the two **P** matrices. A subset of three dimensions was chosen because this is the maximum number of dimensions that can be retained for a comparison of seven-dimensional covariance matrices (k less than or equal to n/2; Aguirre et al. 2014). This approach allows a bounded sum for the eigenvalues of **S** ranging from 0 to 3, representing no and complete shared subspace, respectively. We chose to retain as many dimensions in **S** as possible given that we also compute the vector correlations between the leading eigenvectors of the matrices being compared. We used these pairwise approaches to compare the following matrices: 1) The **P** matrix pooled across all 35 populations and the among-population divergence matrix **D**, which is a test of the degree to which populations have diverged in the same set of traits that vary most across individuals within a population 2) the pooled estimates of Marine, Lake and Stream **P**, and finally 3) Lake and Stream **P** for each watershed where statistical support for ι1**P** was found based on model comparison of mixed models fit separately for each watershed. Finally, because these approaches say little about the overall size of matrices, we also computed evolvability (Bolstad et al. 2014) and conditional evolvability statistics (Hansen and Houle 2008) using random skewers (Cheverud 1996, Cheverud and Marroig 2007). These statistics represent the expected evolutionary response in the direction of selection, where random selection gradients are sampled in multivariate traits space; the distribution of evolvability statistics from such an approach provides an indication of the degree of variational constraints imposed by **P** and can be compared across **P**-matrices.

As a second approach to understanding variation in **P**, we performed an analysis of the 4th order covariance tensor **Λ_P_**, which summarizes variances and covariances of the elements of **P** among populations (Melo et al. 2015). For seven-dimensional **P** matrices, **Λ_P_** can be decomposed into 28 dimensions with corresponding ‘eigentensors’ and their eigenvalues. These eigentensors can be interpreted like PC vectors, except that each eigentensor is a 7x7 dimensional matrix (for 7 traits). Each eigentensor can itself be decomposed into its eigenvectors and eigenvalues (which can be negative); the leading eigenvector from the first eigentensor is the linear combination of traits along which phenotypic variation has changed the most among populations. Hine et al. (2009) and Aguirre et al. (2014) provide an overview of this eigentensor approach. To summarize variation in **P** captured by this covariance tensor, we plotted the first two eigentensors and also calculated the vector correlation between **Λ_P_max,_ d**_max_, and **p**_max_ (calculated from the average of all population-level **P** matrices; nearly quantitatively-identical results are obtained using the pooled estimate of **P** from the ‘common’ model). These vector correlations indicate the degree to which the major axis of variation in **P** does or does not align with the major axis of variation among populations and individuals, respectively.

### Testing predictions for the evolution of **P**

Finally, because lake-stream population pairs represent replicated cases of recent divergence into disparate environments, we can directly test microevolutionary predictions for evolution of **P** under directional selection. Specifically, because the evolution of **P** depends on evolution of **G** and following Phillips and Arnold (1989) and Lande (Lande 1980, 1984), we can describe the within-generation change in genetic variance due to directional selection as

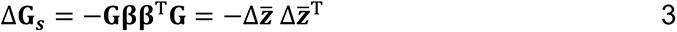

where 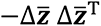 is a matrix of unit rank describing loss of variance in the direction of evolution of trait means under directional selection. Equation 3 describes within-generation changes due to selection, yet these changes are expected to at least partially accrue to the next generation under realistic distributions of allelic affects (Barton 2022). Given the above and the realization that Δ**P**_**S**_ ∝ Δ**G**_**S**_, we expect the matrix Δ**P =** (**P**_stream_ – **P**_lake_) across lake-stream habitats to be aligned with 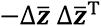 within a given freshwater watershed. Expanding out to consider the set of 16 replicate freshwater watersheds, we expect the covariance tensors Σ_ΔP_ and 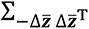 to be aligned if adaptive divergence in trait means drives variation in phenotypic covariance (Hine et al. 2009). Thus, we compared leading eigentensors of these two covariance tensors Σ_ΔP_ and 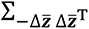, and generated a null distribution for assessment of statistical significance by sampling random Δ**z̄** vectors.

Complete R script and data to reproduce all analyses and figures are deposited on Zenodo (https://doi.org/10.5281/zenodo.8099875).

## RESULTS

Model rankings indicate statistical support for variation in **P** at all levels of analysis; the highest ranked model includes variation in **P** matrices among all 35 populations (Table 1). The same overall conclusions were obtained in an analysis excluding the three marine populations. Thus the model rankings indicate significant variation in phenotypic covariance structure between marine and freshwater, across freshwater watersheds, and at the lowest level of replication, the watershed x habitat population. Pooled estimates of **P** from each of lake, stream, and marine habitat are plotted in Figure 1. Although we find statistical support for divergence in these pooled **P** matrices, **P** was generally similar between lake, stream and marine environments, both in terms of Krzanowski subspace comparisons (Figure S1) and evolvability metrics, although there was some evidence of greater evolvability in the pooled estimate of **P** from marine habitat (Figure S2). Consistent with this finding, we found evidence of increased size of marine **P** in a linear model with the trace of population-specific **P** as the response variable and habitat type (lake, stream, or marine) as a fixed effect (t = 2.999, *p* = 0.00521, df = 32; we found no evidence of a difference between the average size of lake and stream **P** in the same analysis, t = -.278, *p* = 0.78).

**Figure 1.**
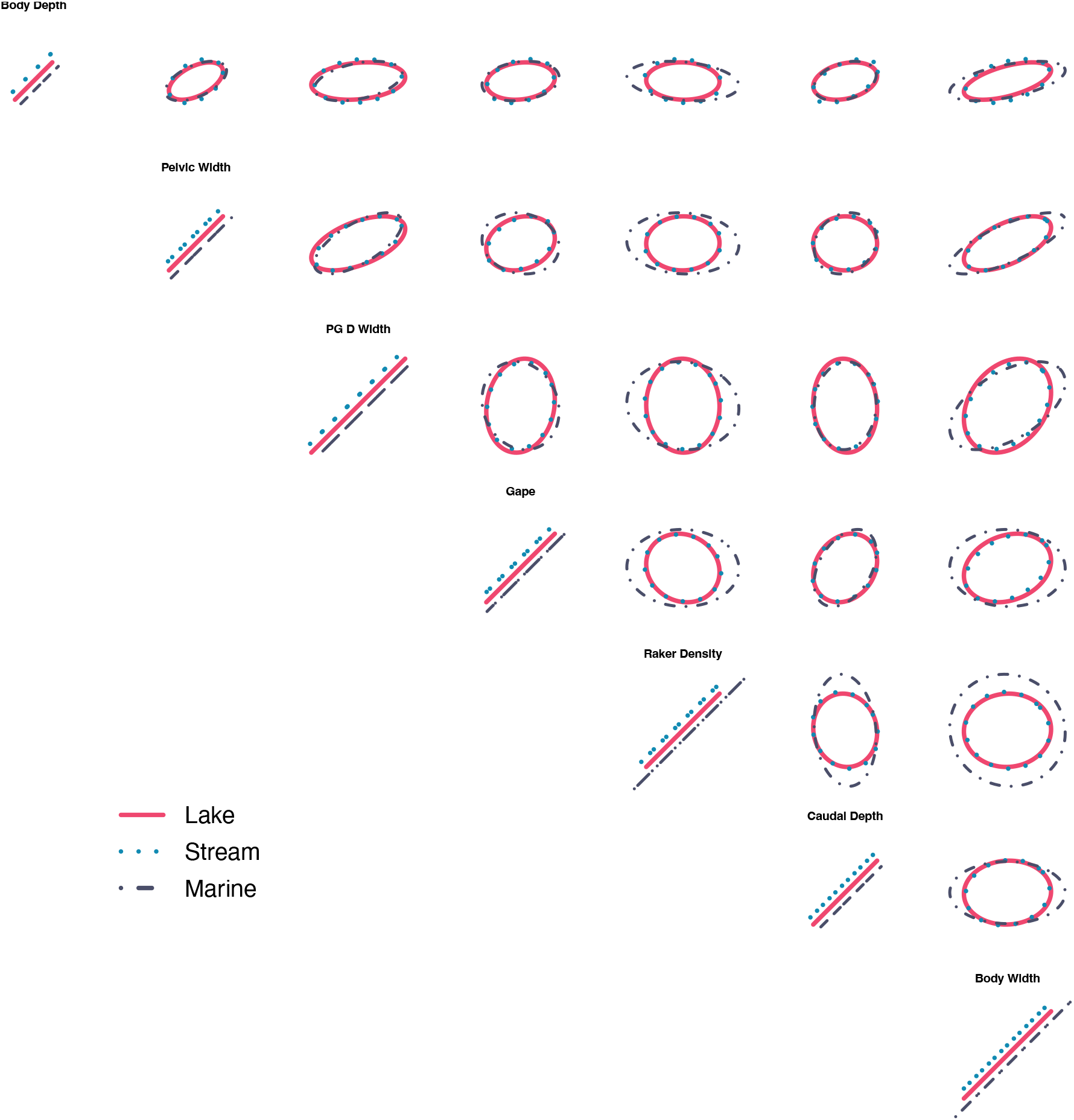
Estimates of seven-dimensional P matrices across marine, and lake and stream freshwater environments. Shown are the posterior modes from the ‘Habitat- specific (3 **P**)’ model.

**Table 1.** Model comparison of multi-response mixed

| **Model** | **P matrices** | **DIC** | **ΔDIC** |
| --- | --- | --- | --- |
| Population-specific | 35 | -24338.24 | 0 |
| Watershed-specific | 19 | -24305.26 | 33 |
| Habitat-specific | 3 | -24235.52 | 102.7 |
| Fresh vs Marine | 2 | -24192.52 | 145.7 |
| Common | 1 | -24101.49 | 236.8 |

We found strong alignment between the **D** matrix describing covariance in mean trait values among populations and the pooled estimate of within-population variation **P** from our ‘common’ model, both in terms of association of leading eigenvectors (vector correlation = 0.91, 95% credible interval for |*r*|= 0.83 – 0.97; Figure 2) and shared Krzanowski subspace (sum of eigenvalues of **S** = 2.58, 95% credible interval = 1.99 – 2.85; Figure 2). This indicates the primary axes of phenotypic variance are nearly completely shared both among individuals within populations, as well as among population means. The lake and stream freshwater **P** matrices tended to be more similar to each other than to the marine **P** matrix in our estimates of **P** from the ‘habitat specific (3 **P**)’ model (Figure 1, S1). However, most of the variation in **P** occurs across freshwater populations and watersheds, as illustrated by the first two eigentensors of Λ**_P_** (Figure 3), indicating that divergence across freshwater populations in both lake and stream environments is unique from marine variation in **P**.

**Figure 2.**
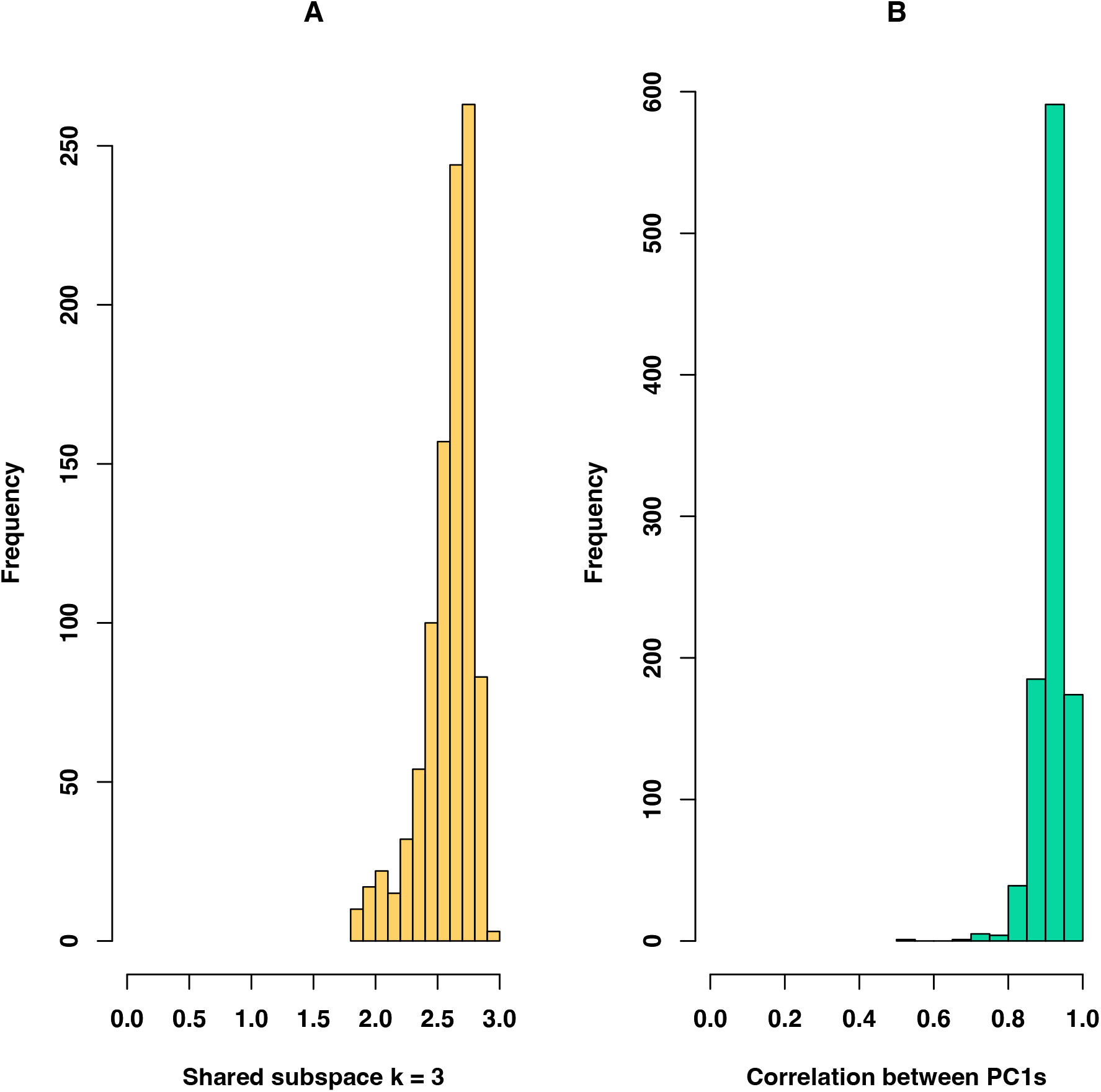
Within and between population variation align. Panel A shows the Krzanowski shared subspace for three dimensions between **D**, the matrix describing variation and covariation in trait means across populations, and **P,** here the pooled estimate from the ‘Common’ model, sampled across the posterior distribution. This is essentially the proportion of the subspace captured by the 3 largest dimensions, with a value of 3 indicating very high similarity and 0 indicating unrelated matrix structure. Panel B shows the vector correlation between PC1 of **D** (**d**_max_)and PC1 of P (**p**_max_), sampled across the posterior.

**Figure 3.**
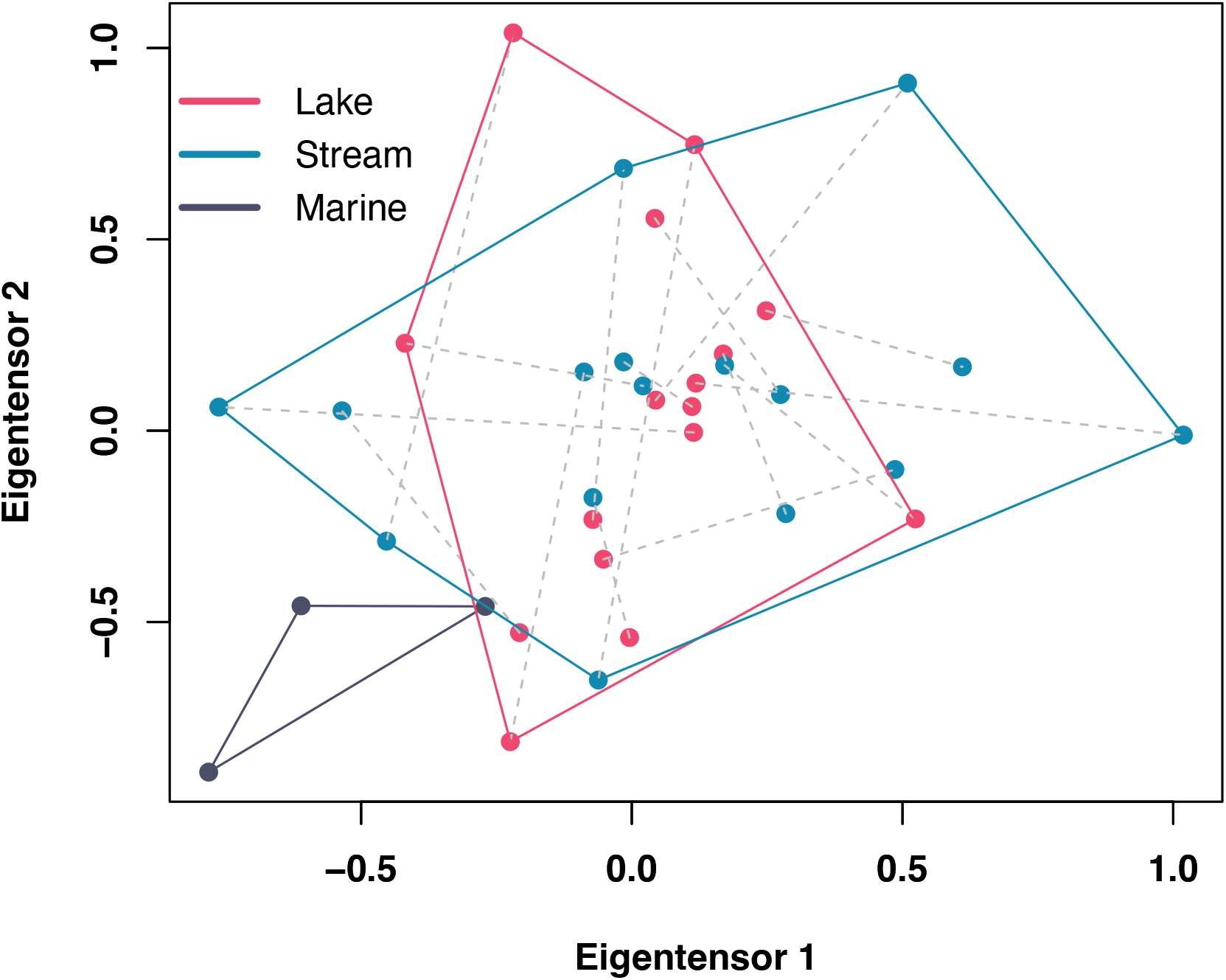
The first two eigentensors of the fourth order covariance tensor **Λ_p_**, which summarizes variation and covariation in elements of **P** across populations. These two tensors capture approximately 50% of the variation in **P**. Points show population- specific estimates from the ‘Population-specific’ model; colors and convex hulls show habitat types, lake and stream populations from the same watershed are connected by dashed grey lines.

We found no evidence of a strong association between Λ**_P_**__max_, the major axis of variation in **P** among populations, and**, d**_max_, the leading eigenvectors of the **D** matrix (vector correlation = 0.32, 95% credible interval = -0.57 – 0.73), indicating that although phenotypic variation among individuals within populations tends to correspond to the pattern of divergence among populations (Figure 2), variation among populations in phenotypic variances and covariances do not occur in these same combinations of traits. Moreover, Λ**_P_**__max_ explained relatively little variation in population means summarized by the **D** matrix (10.8%, 95% credible interval 2.1- 41.8%), and projecting **D** onto **Λ_P_** indicates that this the directions of divergence among populations doesn’t explain much of the population-to-population variation in **P** (only 0.012% of the variance in **P)**. To appreciate these findings, we can consider the loadings for the principle components of phenotypic variation for the traits that load most and least strongly on **p**_max_. Figures 4A and 4C show the two traits that load most strongly on PC1 of the average **P** matrix (taken across all freshwater and marine populations), body width and pelvic girdle diamond width.

**Figure 4.**
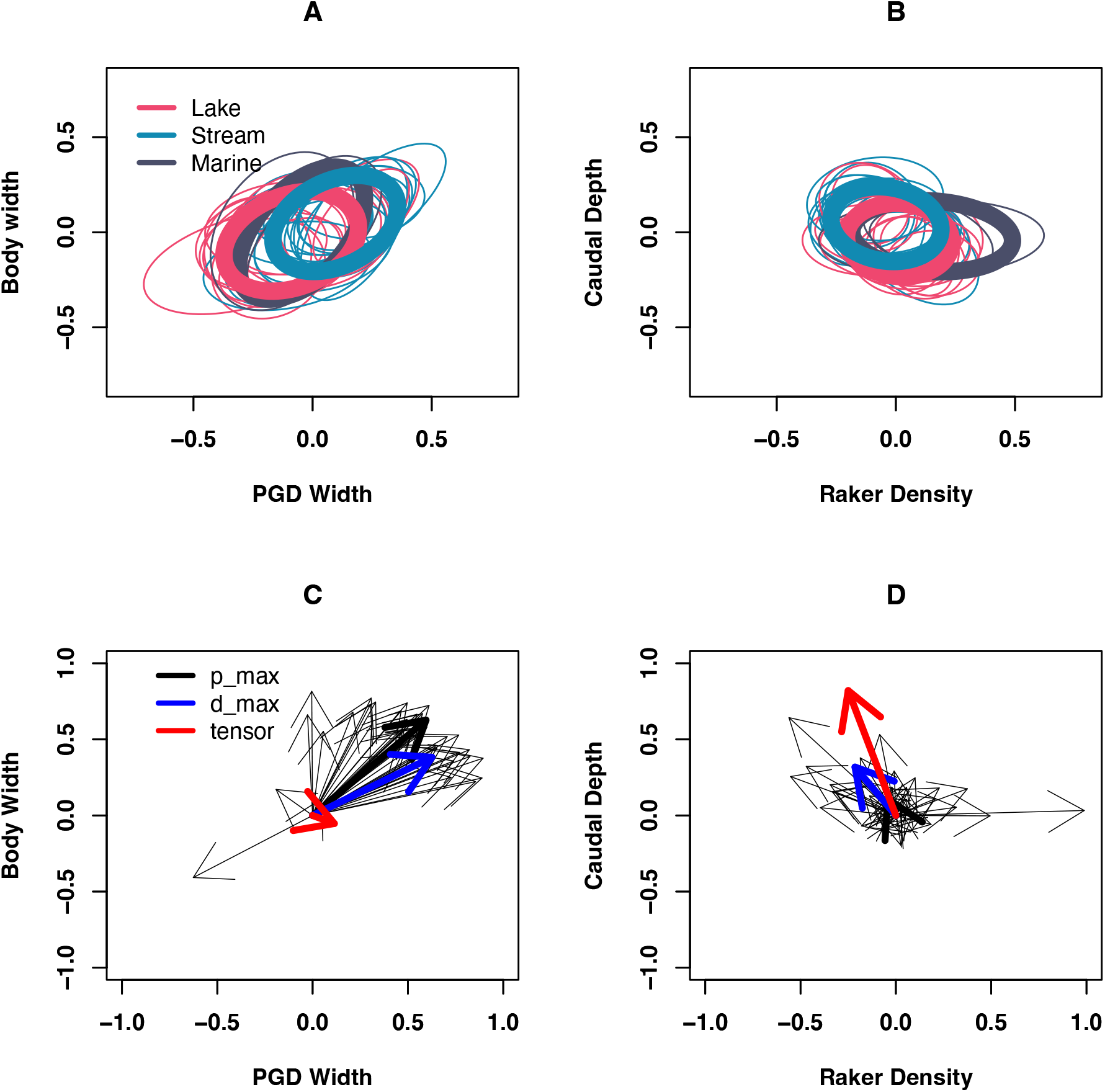
Variation in P matrices among populations. Panels A and B show estimates (posterior modes) of P (plotted for two different traits in each panel) for each of the 35 populations sampled, as well as the pooled estimates for lake, stream, and marine environments shown by thick ellipses. Panels C and D show variation in the orientation of major axes of variation across populations for the same sets of traits. Panel C shows the loadings for the two traits that load most strongly on PC 1 from the average **P** matrix, or **p**_max_; the average is shown by the thick black arrow and thin black arrows show population specific estimates. The blue arrow shows PC1 from the **D** matrix, or **d**_max_; this is the direction of maximum variation among population means. The red arrow shows PC1 from the first eigentensor of **Λ_p_**, which describes the combination of traits that vary most in their degree of (co)variance among populations. Panel D shows the same but for raker density and caudal depth, two traits that do not load strongly on average on **p**_max_. As panel C shows, these two traits, body width and pelvic girdle diamond width, tend to be tightly correlated within and across populations, with relatively little variation in the degree of this correlation. Alternatively, caudal depth and raker density show substantial variation in their (co)variance among populations, shown in Panel D.

These two traits tend to be strongly correlated (see also Figure 1). Figure 4C shows not only alignment of average **P** and the **D** matrix, but also relatively low variation in covariance across populations, as illustrated by the low loadings of these traits on Σ_P___max_, PC 1 of the first eigentensor of Σ_P_. In contrast, Figures 4B and 4D show the two traits that load least strongly on this component, gill raker density and caudal depth. These two traits have low correlations with each other (Figure 4B and Figure 1). Although PC1 of **D** and average **P** are aligned here despite low loadings, there is substantial population variation in the orientation and size of **P**, reflected in the high loadings of these two traits on PC1 of the first eigentensor of Σ_P_ (Figure 4D). Thus, these data indicate patterns of strongly conserved covariance structure within and among populations in some suites of traits (Figure 4A, 4C), along with patterns of divergence in covariance of ecologically important traits that are less strongly integrated (Figure 4B, 4D).

In our test of quantitative genetic predictions for evolution of **P**, we found a striking relationship between variation in lake-stream 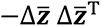, the unit-rank matrix describing expected change in variance due to directional selection, and lake-stream Δ**P**. That is, the correlation between the leading eigenvector of the first eigentensor of Σ_ΔP_ and 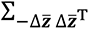 is near one and highly significant (vector correlation = 0.96, *p* < 0.0001; Figure 5), indicating that variation in Δ**P** among freshwater watersheds was greatest in the direction of trait space where Δ**z̄** also varied most. Moreover, 33% of the variation in Δ**P** can be explained by variation in 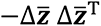 (*p* < 0.0001; Figure 5). Thus, variation across watersheds in the change in **P** between lake and stream environments is predicted by variation in evolution of mean trait values across these watersheds. This is consistent with multivariate quantitative genetic expectations (Lande 1980, 1984, Phillips and Arnold 1989), and indicates that variation in evolutionary change of **P** between lake and stream environments matches variation in evolutionary change in means between these environments across watersheds. Thus, among-watershed differences in Δ**P** are predictable by among-watershed differences in Δ**z̄**, although we note that this association cannot rule out the possibility that Δ**z̄** was influenced by Δ**P** rather than viceversa.

**Figure 5.**
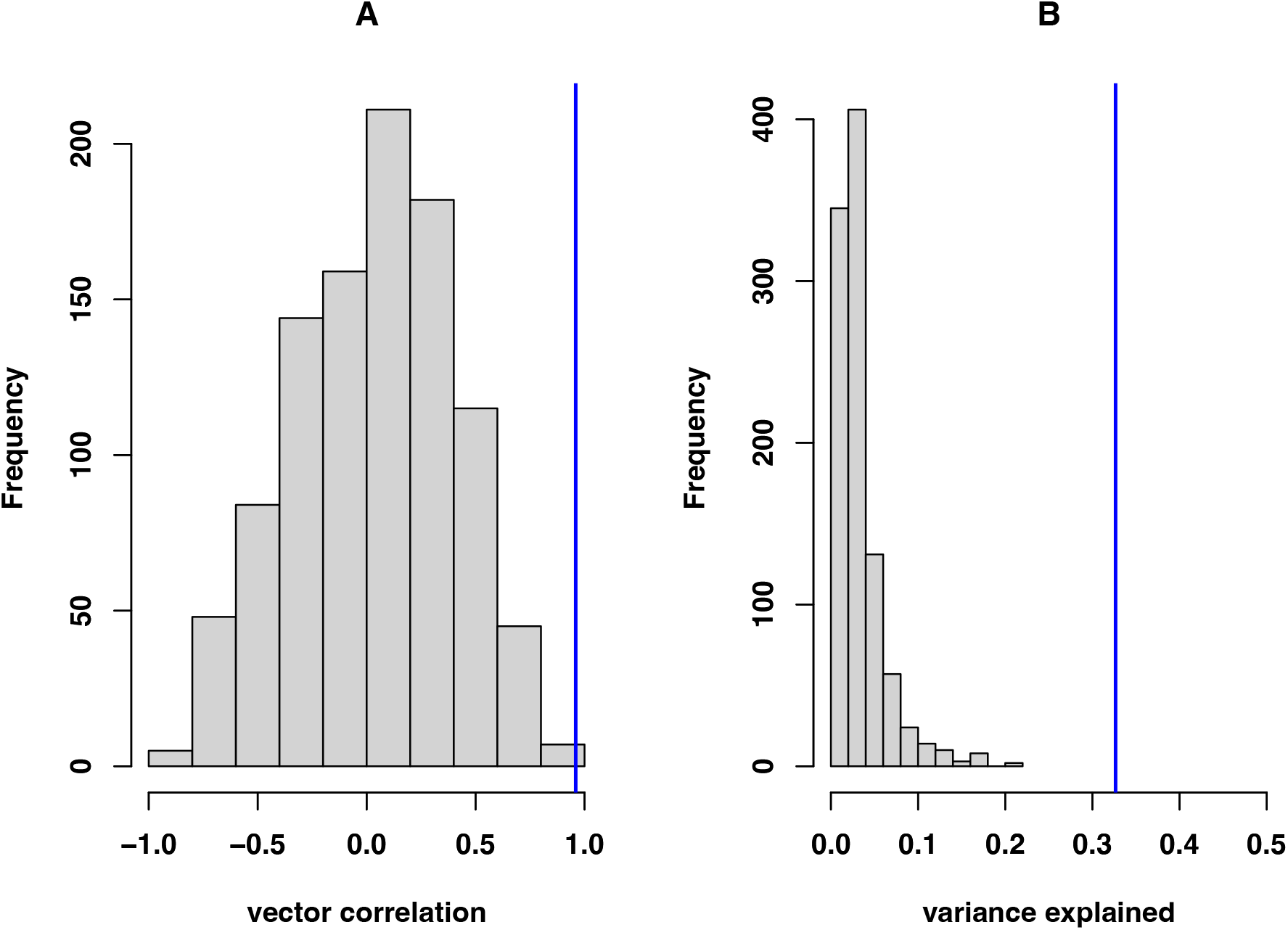
Variation in ΔP across freshwater watersheds is explained by variation in Δ**z̄**. Panel A shows the vector correlation between the leading eigenvector of the first eigentensor of Σ_ΔP_ and 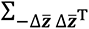 (observed, blue; grey, null); that is, the single linear combination of traits that explains the most variation in each of these two matrices is nearly identical. Panel B shows the variance in ΔP explained by the first eigentensor of 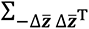 (observed, blue; grey, null).

We next explored within-watershed predictions, focusing only on watersheds where there was statistical support for evolution of **P** between lake and stream environments (to avoid interpreting noise). We identified 4 watersheds where there was robust support for lake-stream divergence in **P**, based on ΔDIC > 7 (Table 2). For each of these four watersheds, we found significant change in phenotypic variance between lake and stream populations in the direction of evolution of trait means, as captured by the projection 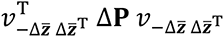, where 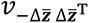 is the eigenvector associated with the most negative eigenvalue of the matrix 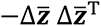 (Figure 6; Beaver, Bayesian *p* = 0.046; Boot, Bayesian *p* = 0.034; Moore, Bayesian *p* = 0.001; Roberts, Bayesian *p* = 0.019), and these effects are visible in the form of a change in the 95% bivariate (co)variance ellipses for the two traits that load most strongly on Δ**z̄** for each watershed (Figure 6). Thus we find significant evolution of variance in the combination of traits captured by Δ**z̄** in populations where there is strong statistical support for Δ**P**. We note that although theory predicts a reduction of variance, we cannot assess this because we do not know the polarity of lake-stream change. For reference, we found no consistent patterns of change in evolvability between lake and stream **P** in these four watersheds (Figure S4) or variance evolution along other eigenvectors of 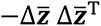. (Figure S5), a dimension not expected to be associated with variance evolution under theoretical expectations (Phillips and Arnold 1989).

**Figure 6.**
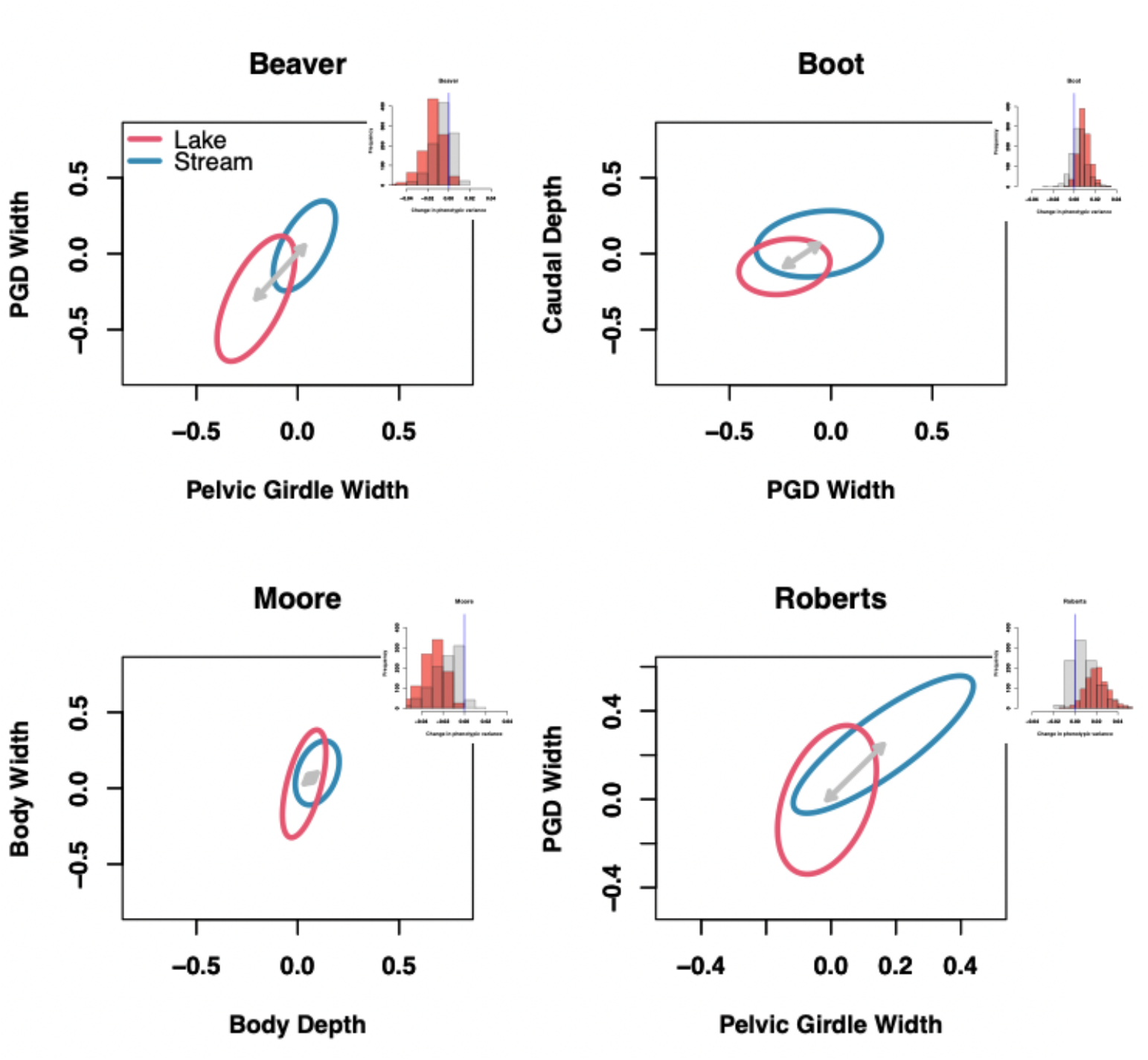
Evolution of phenotypic variance along predicted direction of Δ**z̄**, from watersheds with robust statistical support for Δ**P**. Each panel illustrates change in **P** for the two traits that contribute most to Δ**z̄** in the watershed in question; grey arrows show Δ**z̄**. Inset red histograms show 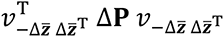, where 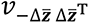 is the eigenvector associated with the most negative eigenvalue of the matrix −Δ**z̄** Δ**z̄**., sampled from the posterior distribution with bootstrapped Δ**z̄**. Light grey shows samples generated using random samples from the entire data set rather than population lake or stream habitats. These inset panels thus show the observed change in variance in the direction of Δ**z̄**.

**Table 2.**
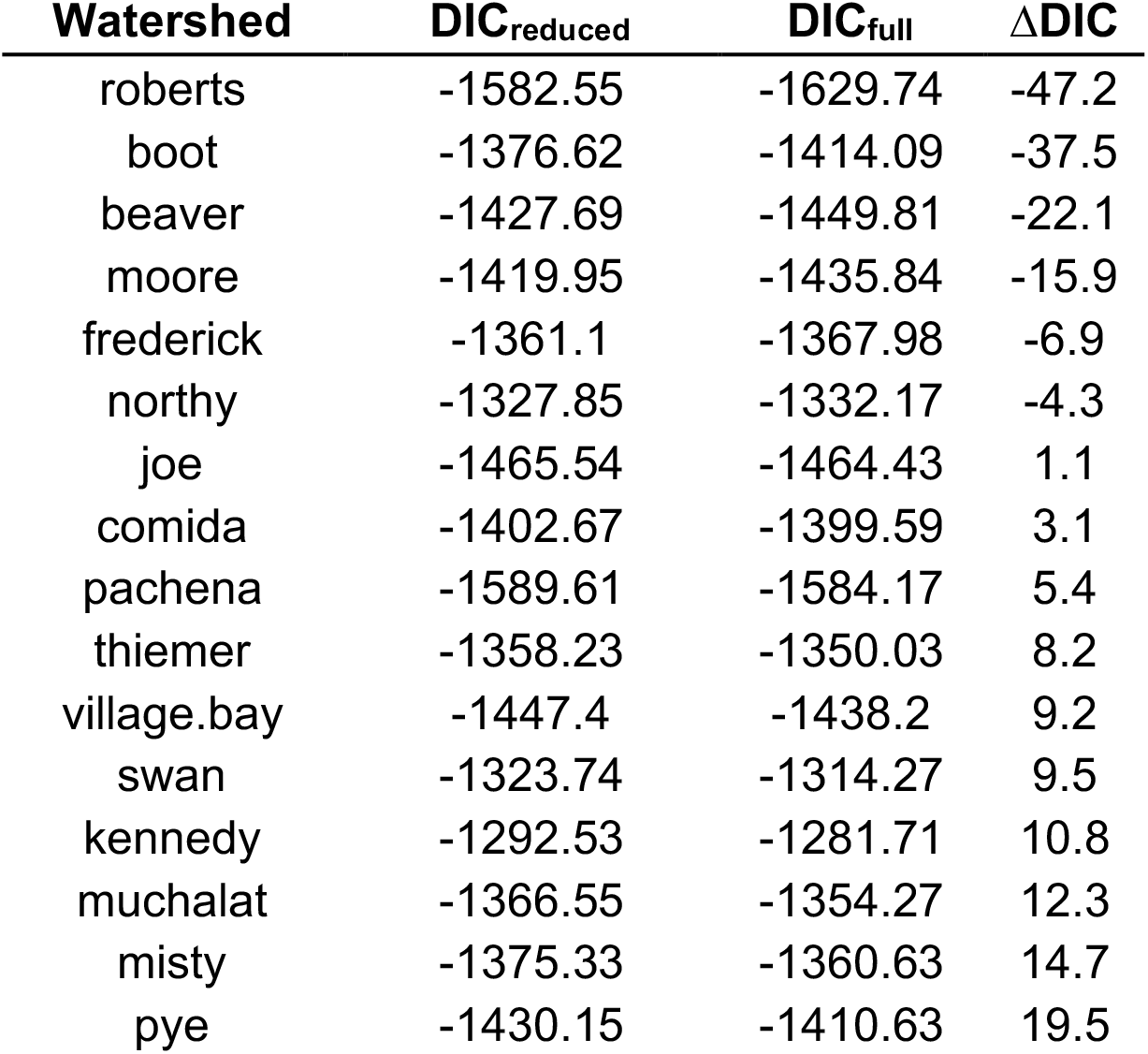
Model comparison to assess lake-stream differences in P for each watershed, from nested multiresponse mixed modes fit separately for each watershed

## DISCUSSION

We analyzed multivariate phenotypic variation across a set of 35 populations of threespine stickleback (*Gasterosteus aculeatus*) to understand how phenotypic covariance has evolved during replicated adaptation to freshwater environments (Stuart et al. 2017). We found evidence of significant divergence in **P** matrices across populations against a background of what is, largely, shared covariance structure. The major axis of phenotypic variance is strongly aligned across populations and is further aligned with divergence in mean trait values across populations. Evolution of phenotypic variance, in contrast, has occurred in suites of traits that tend to exhibit low correlations with other traits (low integration). Simple microevolutionary predictions for evolution of variance are surprisingly successful in predicting change in **P** matrices across lake and stream environments within watersheds, indicating that watershed-specific changes in the covariance of ecologically important traits, such as gill rakers and caudal depth, have occurred during adaptation to disparate freshwater environments.

By far, most variation in phenotypic (co)variance occurred between freshwater and marine habitats and among freshwater watersheds; we found less evidence of consistent differences between lake and stream **P** matrices. This suggests that variation in **P** was largely idiosyncratic, and non-parallel across the replicated instances of adaption to lake and stream environments similar to the patterns of non-parallelism in means observed in the same set of populations (Stuart et al. 2017, De Lisle and Bolnick 2020). This is illustrated in our analysis of the covariance tensor describing among-population variation in **P**, where dispersion along the first two eigentensors illustrates the substantial variation in **P** and lack of consistent differences between lakes and streams. This result is also demonstrated in our comparison of candidate models where the most dramatic drops in information criteria were observed when including separate **P** matrices for freshwater versus marine, and subsequently for separate watersheds. Marine populations exhibited less variation in **P** and exhibited tighter covariance in traits related to body shape and substantially more variance in gill raker density, relative to freshwater populations. However, it is notable that despite these disparities in **P**, our pooled estimates of marine, lake, and stream **P** matrices were in large part very similar in structure.

Although we found little evidence of consistent parallel evolution of **P** among lakes and among streams, within-watershed differences in lake and stream **P** matrices were in some cases substantial. Moreover, these differences lake and stream **P** within watersheds were predictable based on simple quantitative genetic predictions for within-generation change in variance. More precisely, variation in lake-stream Δ**P** across watersheds was predictable based on variation in lake- stream Δ**z̄**, indicating that seemingly-idiosyncratic variation in **P** is in fact explainable based on non-parallel evolution of population means across watersheds. This variation in Δ**P** indicates that changes in phenotypic covariance structure occur in the same suites of traits that show greatest divergence in mean value across lake and stream habitats within specific watersheds. Such within-watershed predictability of evolution of **P** does not scale up to generate consistent differences in average lake and stream **P** matrices across watersheds, however, likely because of the substantial variation in the observed directions of multivariate trait differences between lake and stream environments across watersheds. That is, in analyses of trait means, Stuart et al (2017) found little evidence of repeatable lake-stream divergence. Applying another analytical approach to the same data, De Lisle and Bolnick (2020) supported the conclusion that divergence across lake-stream habitat is more complex than can be described by a single dimension of parallel evolution (i.e., the lake-stream axis). Our study suggests that this among-watershed variation in the direction of multivariate evolution has driven subsequent evolution of variation in phenotypic covariance structure across this suite of populations—a continuum of (non)parallelism in P-matrices. Noteworthy in this regard is that our analysis is based on a subset of traits that have contributed most to parallel evolution of multivariate mean phenotypes in this suite of populations (De Lisle and Bolnick 2020); even with this focus on the traits contributing most to parallel evolution, we found substantial variation across watersheds in how the **P** matrix changed between lake and stream environments.

We have focused on understanding variation in patterns of phenotypic (**P**), rather than genotypic (**G**), variance. A justification for this is that selection acts on phenotypic variance, and that evolution of **P** must be shaped by the same phenomenon that may affect evolution of **G** (Cheverud 1988). Nonetheless, a major caveat is that we cannot account for changes in **P** induced by variation in environmental (co)variance across habitats and populations. Indeed, reanalysis of a previous study of some of the same populations (Oke et al. 2016) indicates that **P** matrix estimates can differ between wild-caught fish and those reared in the laboratory, although generally share similar structure (see supplemental analysis), suggesting an important contribution of environmental effects on **P** in stickleback. Future studies examining the contribution of genotype x environment interaction to patterns of phenotypic variation in this radiation would be informative.

Schluter (1996) proposed, and ample evidence has accrued (Blows and Higgie 2003, McGuigan et al. 2005, Chenoweth et al. 2010, Punzalan and Rowe 2016, McGlothlin et al. 2018), that bias in response to selection produced by multivariate genetic constraints may generate a signature of association between major axes of genetic variance and mean population divergence. More generally, there is a long standing empirical finding that, across traits within a study system, measures of within population variation coincide with among-lineage divergence (Kluge and Kerfoot 1973, Bolstad et al. 2014, Houle et al. 2017). An underappreciated (although noted by Schluter and others; e.g. Punzalan and Rowe 2016) alternative interpretation of such patterns is that selection shapes evolution of both trait means and genetic variance on timescales relevant to produce a correspondence between the two. Our finding of an association between variation in Δ**z̄** and Δ**P** supports this alternative interpretation of patterns of evolution along genetic lines of least resistance, in that our finding is consistent with selection shaping both standing phenotypic variance and divergence in trait means across lake and stream environments.

Our findings that multivariate variation largely aligns with divergence and concomitantly rapid evolutionary change in variance, is consistent with patterns found in another adaptive radiation. In their analysis of *Anolis* **G** matrix evolution, McGlothlin et al. (McGlothlin et al. 2018, McGlothlin et al. 2022) found that evolution of **G** occurred primarily in the same combinations of traits that varied most among individuals and among populations. This differs from our finding, where we observe most of the divergence in **P** matrices occurs largely orthogonally to ***p*** or ***d*** max, indicating that most evolution of variance occurs in different suites of traits than those that are involved in conserved divergence within and among populations.

However, consistent with our finding, McGlothlin et al (2022) found that predictable features of **G** matrix evolution in Anolis ecomorphs have occurred primarily in suites of traits that, in general, show less integration with other traits. This is consistent with our findings, where predictable features of change in **P** occur in suites of stickleback traits that are not consistently tightly integrated.

Our work paints a nuanced picture of the evolution of phenotypic variation in stickleback. On the one hand, patterns of variation of tightly covarying traits, such as pelvic traits and body width, remain deeply conserved within and across populations. On the other hand, some combinations of traits show rapid change in (co)variance between habitats that is predictable from simple quantitative genetic theory. Thus, our study adds to a growing body of work (Hohenlohe and Arnold 2008, Punzalan and Rowe 2016, McGlothlin et al. 2018, McGlothlin et al. 2022) that indicates a potential role for selection in contributing to apparent correspondence between within-population variation and among population divergence. Our work highlights how both conservation and rapid divergence of multivariate variation can both contribute to patterns of divergence in adaptive radiation.

## Acknowledgements

We are grateful to David Houle, Kjetil Voje, and Masahito Tsuboi for discussion and feedback. S.D. was supported by grants from the Swedish Research Council (2019- 03706), Formas (2021-01096), and the Crafoord Foundation (20220602). Y.E.S was supported by grants from the US National Science Foundation (DEB-1144773 ; DEB-1456462; EAR-2145830).

## Glossary of Terms

| term | dimensions* | definition |
| --- | --- | --- |
| **P** | $k \times k$ | Phenotypic variance-covariance matrix, summarizing variation in trait expression among individuals |
| **G** | $k \times k$ | Genotypic variance-covariance matrix, summarizing variation in additive genetic effects among individuals |
| **D** | $k \times k$ | Among population divergence matrix, a variance-covariance matrix summarizing variation in trait means across populations or lineages. |
| $\Delta \bar{z}$ | $k$ | A vector of evolutionary change in trait means between two populations, in this case, between lake and stream habitats within watersheds |
| $\Delta \mathbf{P}$ | $k \times k$ | A matrix describing change in phenotypic (co)variance between lake and stream habitats, taken as $\mathbf{P}_{\text{stream}} - \mathbf{P}_{\text{lake}}$ |
| $-\Delta \bar{z} \Delta \bar{z}^T$ | $k \times k$ | A unit-rank matrix describing the expected change in **G** or **P** due to directional selection |
| $\mathbf{d}_{\text{max}}$ | $k$ | Leading eigenvector (PC1) of **D**, a vector describing the combination of traits that vary most in mean value among populations |
| $\mathbf{p}_{\text{max}}$ | $k$ | Leading eigenvector (PC1) of **P**, a vector describing the combination of traits that vary most among individuals |
| $\boldsymbol{\beta}$ | $k$ | Vector of directional selection gradients, describing selection on each trait |
| $\Sigma_{\mathbf{P}}$ | $k \times k \times k \times k$ | Fourth order covariance tensor, describing variance and covariance in elements of **P** among populations |
| $\Sigma_{\Delta \mathbf{P}}$ | $k \times k \times k \times k$ | Fourth order covariance tensor, describing variance and covariance in elements of $\Delta \mathbf{P}$ among freshwater watersheds |
| $\Sigma_{-\Delta \bar{z} \Delta \bar{z}^T}$ | $k \times k \times k \times k$ | Fourth order covariance tensor, describing variance and covariance in elements of $-\Delta \bar{z} \Delta \bar{z}^T$ among freshwater watersheds |

\* where $k$ is the number of traits, in this case seven.

## Supporting Information

**Table S1.**
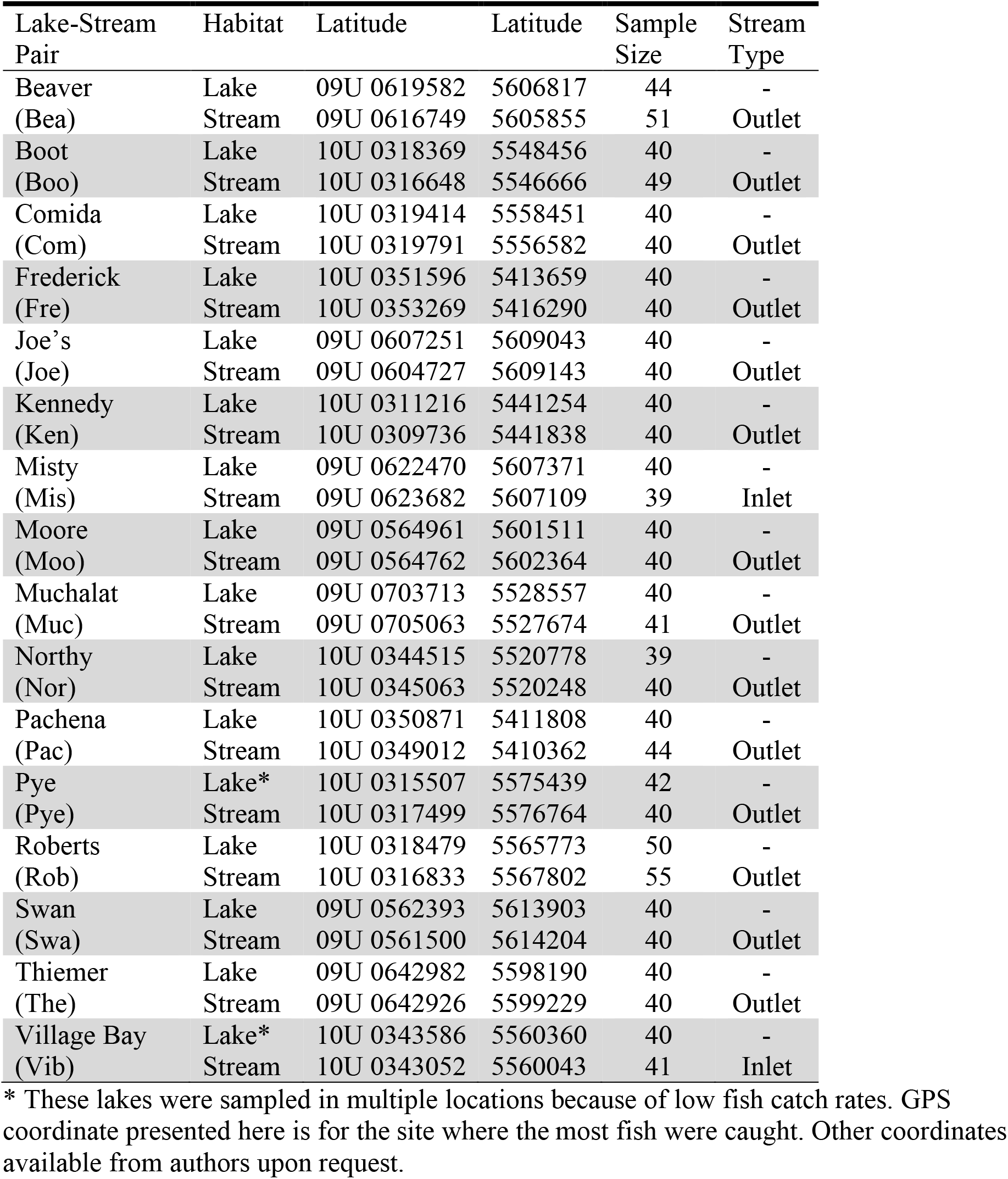
Collection localities and sample sizes. Latitude and longitude reported in UTM units. 40 fish were sampled from each of Campbell Marine (CamM) and Cluxewe Marine (ClxM). 48 fish were sampled from Rupert Marine (RupM).

**Table S2.** Description of 32 linear and meristic morphological measurements plus standard length. Standard length was used to size-correct the 32 linear and meristic measurements and was not included in the rest of the analyses.

| Trait | Method | Description |
| --- | --- | --- |
| First dorsal spine length | Lateral image | Length, anterior insertion to tip |
| Second dorsal spine length | Lateral image | Length, anterior insertion to tip |
| Dorsal fin length | Lateral image | Length, at insertion points |
| Caudal peduncle depth | Lateral image | Depth, narrowest point of caudal peduncle |
| Anal fin length | Lateral image | Length, at insertion points |
| Pectoral fin insertion length | Lateral image | Length, dorsal to ventral insertion |
| Body depth | Lateral image | Depth, anterior insertion of first dorsal spine to anterior point of pelvic girdle |
| Mouth length | Lateral image | Length, inside corner of mouth to anterior tip of lower jaw |
| Snout length | Lateral image | Length, tip of upper jaw to closest point on the perimeter of the eye |
| Eye diameter | Lateral image | Diameter, from posterior end of snout |
| Head length | Lateral image | Length, tip of upper jaw to posterior most point of the gill operculum |
| Pectoral fin width | Lateral image | Width, measured from distal ends of outer fin rays. |
| Pectoral fin length | Lateral image | Length, longest fin ray |
| Pectoral fin perimeter | Lateral image | Perimeter |
| Buccal cavity length | Ventral image | Length, tip of snout to anterior point of ectocoracoid |
| Gape width | Ventral image | Width, distance between mouth corners |
| Body width pt.1 | Ventral image | Width, jawbone to jawbone, centered at the eyes |
| Body width pt.2 | Ventral image | Width, distance between posterior points of ectocoracoid |
| Pelvic girdle width | Ventral image | Width, between insertions of pelvic spines |
| Pelvic girdle length | Ventral image | Length, anterior-most point of girdle to caudal tip of posterior process |
| Pelvic girdle diamond width, posterior process | Ventral image | Width, widest points of posterior process |
| Pelvic girdle diamond length, posterior process | Ventral image | Length, from anterior-most point to caudal tip |
| Body width pt. | Ventral image | Width, across anterior insertion of anal fin |
| Body width pt.4 | Ventral image | Width, across posterior insertion of anal fin |
| Mean pelvic spine length | Calipers | Insertion to tip, left side and right side |
| Standard length | Calipers | Length, tip of snout to end of caudal peduncle |
| Left side plate number | Count | Included small plates, did not include post-temporal bone, left side |
| Right side plate number | Count | Included small plates, did not include post-temporal bone, right side |
| Right side gill raker number | Count | Counted *in situ*, under a microscope |
| First longest raker length | Raker image | Dissected raker, measured from photograph. |
| Second longest raker length | Raker image | Dissected raker, measured from photograph |
| Third longest raker length | Raker image | Dissected raker, measured from photograph |
| Raker density | Raker image | Dissected raker, length spanned by five longest rakers |

**Figure S1.**
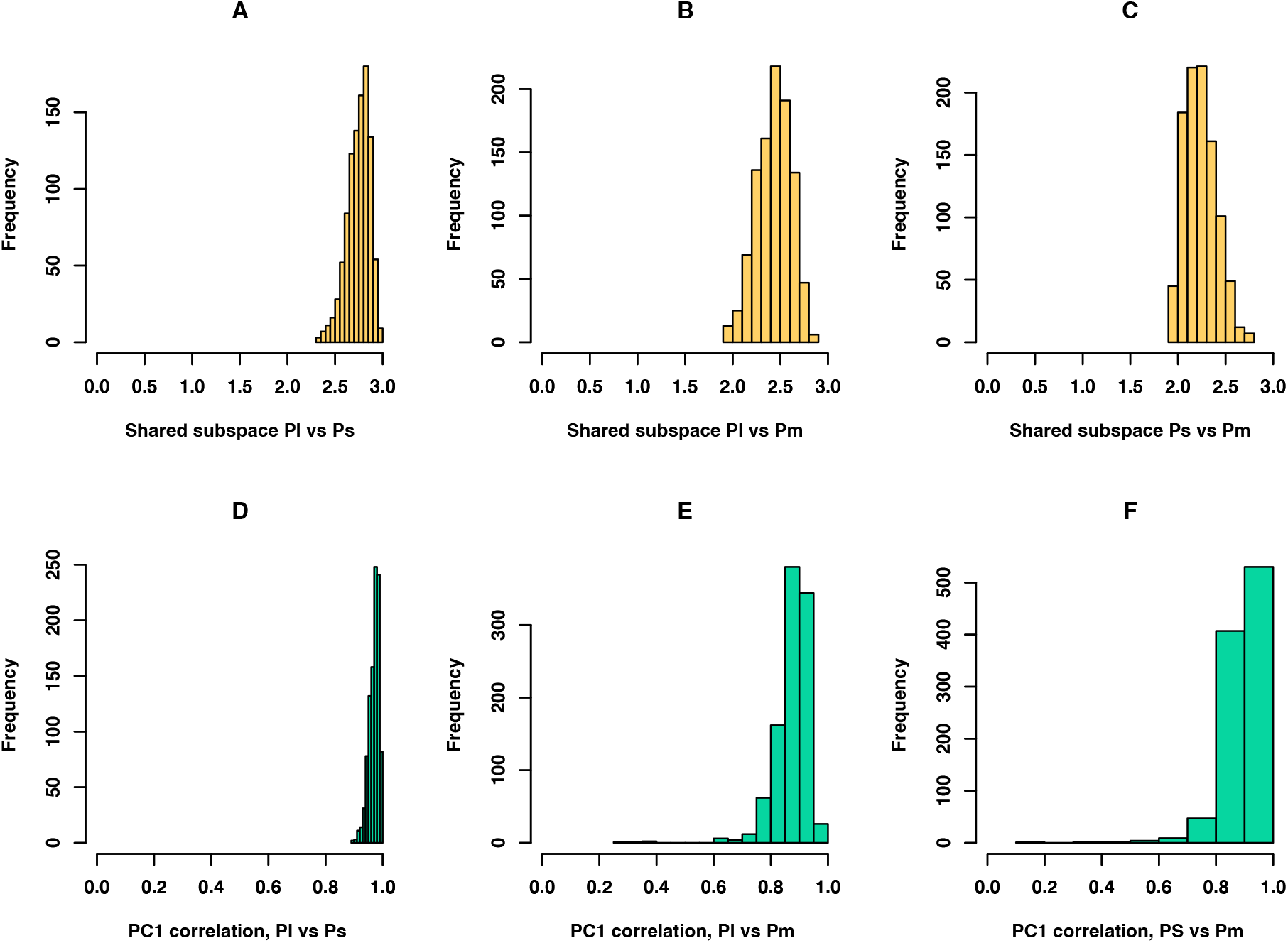
Comparing **P** matrix structure across habitats. Top row shows Krzanowski subspace comparisons for each pairwise contrast between lake and stream (A), lake and marine (B), and stream and marine (C), sampled across the posterior distribution. The bottom panels D-F show the vector correlations between PC 1 of each **P** (**p**_max_) for each of the same pair-wise contrasts. Although all three matrices are quite similar, stream and lake **P** are slightly more similar to each other than they are to marine

**Figure S3.**
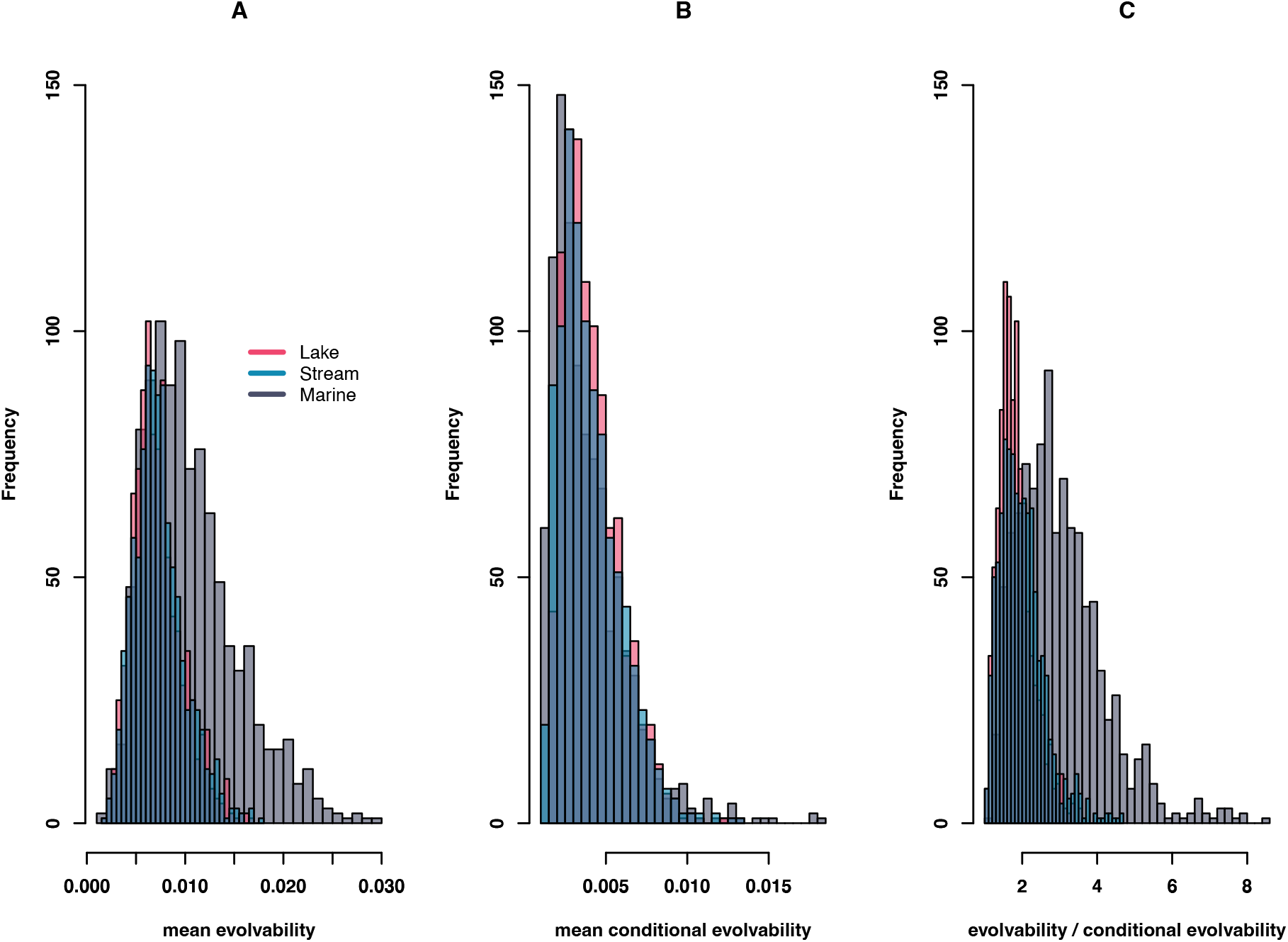
Comparisons of evolvability statistics across Lake, Stream, and Marine. Panel A shows mean evolvability, the expected evolutionary response in the direction of selection; Panel B shows conditional evolvability, which is the expected response to directional selection under the assumption of stabilizing selection on non-selected trait combinations. Panel C shows the ratio between the two. All histograms show samples across the posterior distribution of **P**, generated using random selection vectors in 7 dimensional space.

**Figure S4.**
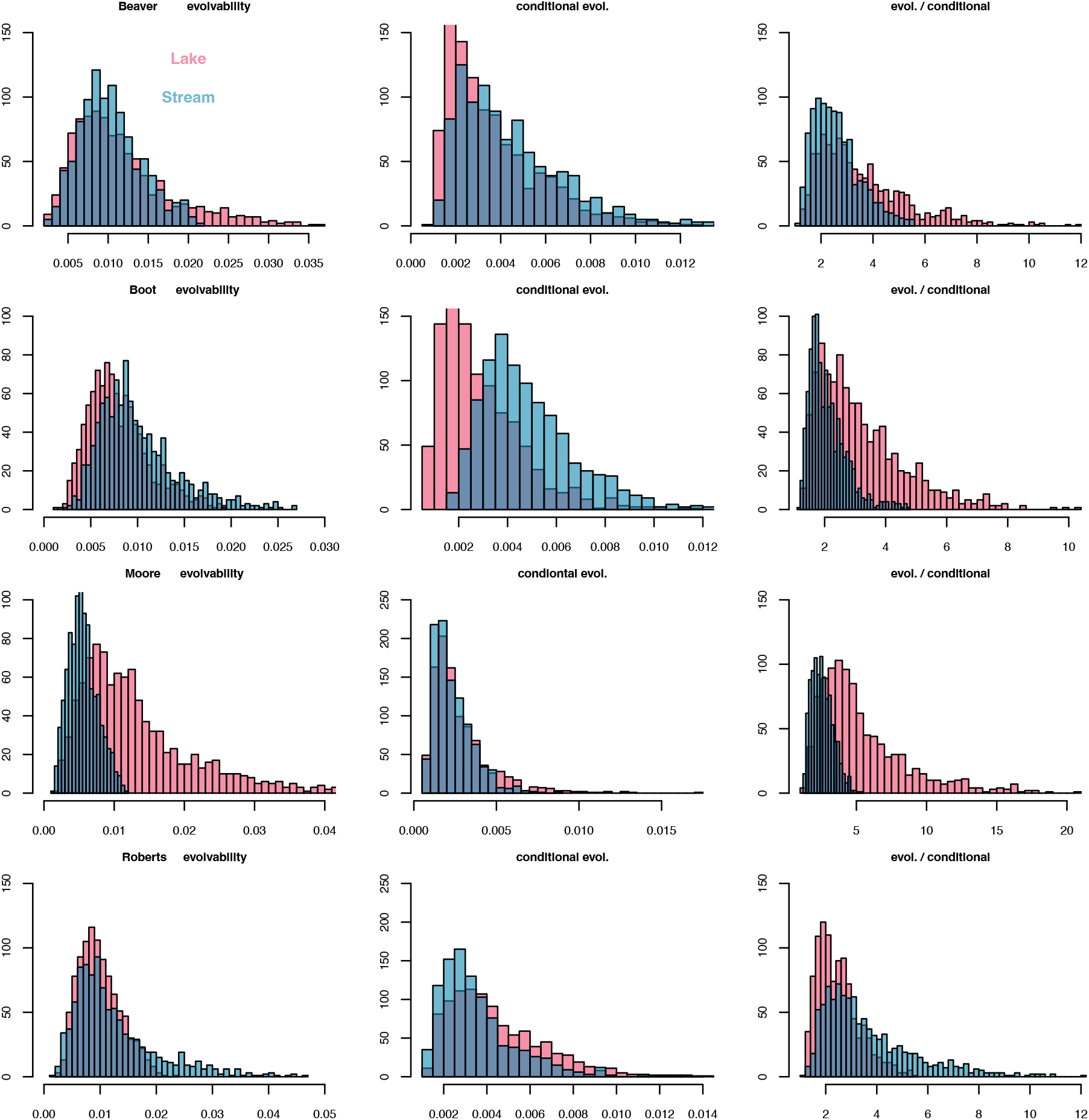
Evolvability across Lake and Stream habitats for watersheds with significant evolution of **P**. Columns from left to right show mean evolvability, mean conditional evolvability, and the ratio between the two sampled across the posterior distribution for each watershed

**Figure S5.**
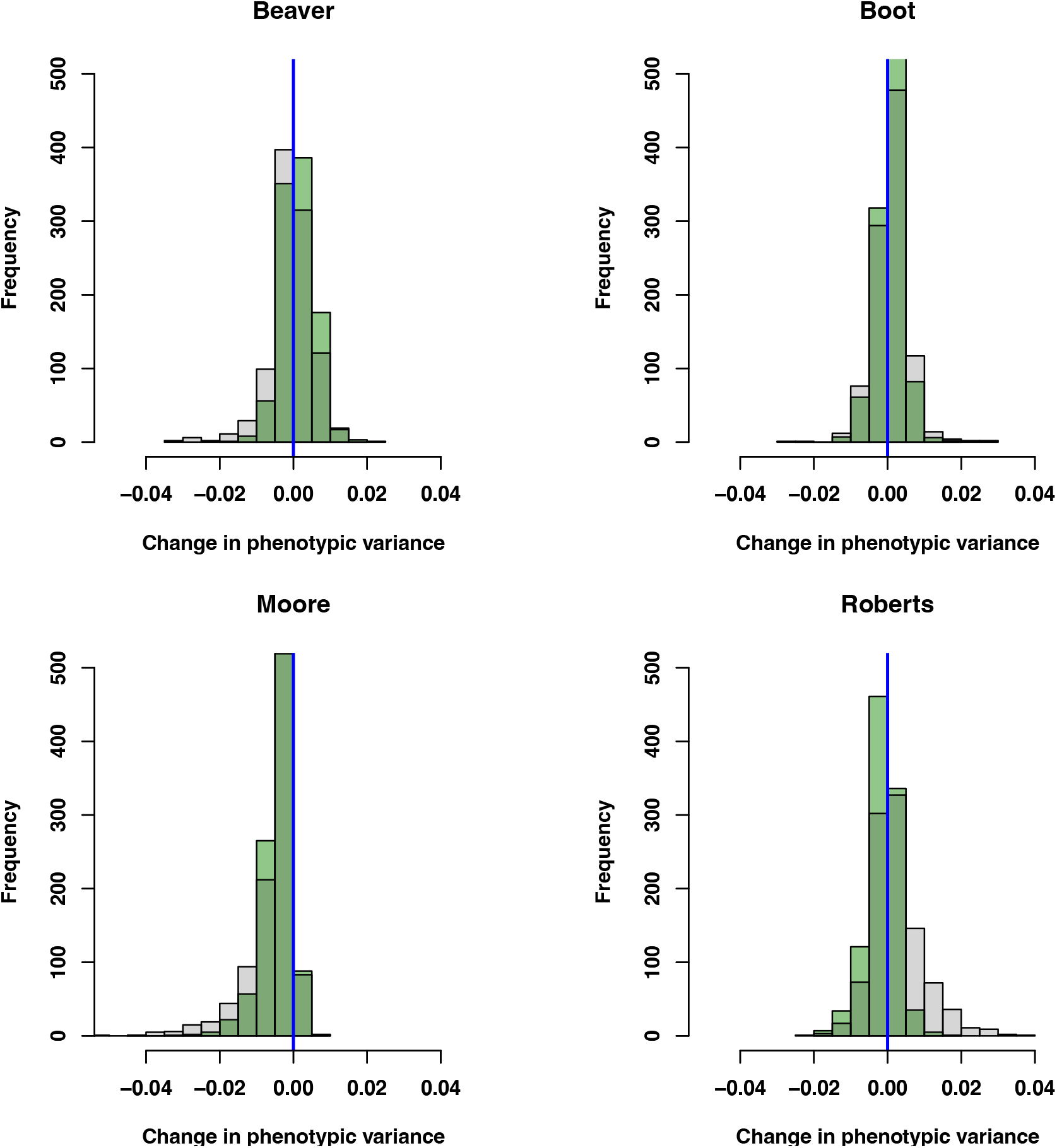
Repeat of the inset panels of figure 6, but illustrating change in variance along most positive eigenvector of −Δ**z̄** Δ**z̄**.; this is a vector associated with a null eigenvalue, for which we don’t predict with variance evolution under our Lande equation prediction. We indeed find that this direction of trait space is not associated with any significant shifts in phenotypic variance. All p > 0.05.

**Supplemental Analysis:** comparison of **P** matrices estimated from lab-reared and wild-caught fish. Here, we reanalyze published data from Oke et al. (2016), who performed a common garden experiment comparing morphological values from wild caught fish to values from lab-reared fish whose genetic background originated from the same population. Oke et al. were interested in understanding the contribution of plasticity to parallelism of means, but their data also allow an assessment of whether estimates of **P** may differ depending on the rearing environment of the fish. They present data from three of the same watersheds studied in the present paper, Misty, Roberts, and Boot. We reanalyzed their data, focusing on seven morphological traits that are closest to the focal traits of our study: Dorsal.fin.length, Caudal.peduncle.depth, Number.of.Rakers, avg.of.Median.GR.length.mm, X1st.dorsal.spine.length, Jaw.length, Pectoral.fin.width. We fit Bayesian multiresponse mixed effects models as described in the main text. Analyzing each of the three freshwater watersheds separately, we fit a model with separate P matrices estimated for lab-reared vs wild-caught fish, and compared this to a reduced model with a single P matrix. We found support for differences in P between lab-reared and wild-caught fish for all three watersheds (Misty, **ΔDIC** = -47; Boot, **ΔDIC** = -20; Roberts, **ΔDIC** = -32), although in general the structure of **P** is similar for both rearing environments (Figure S6).

**Figure S6.**
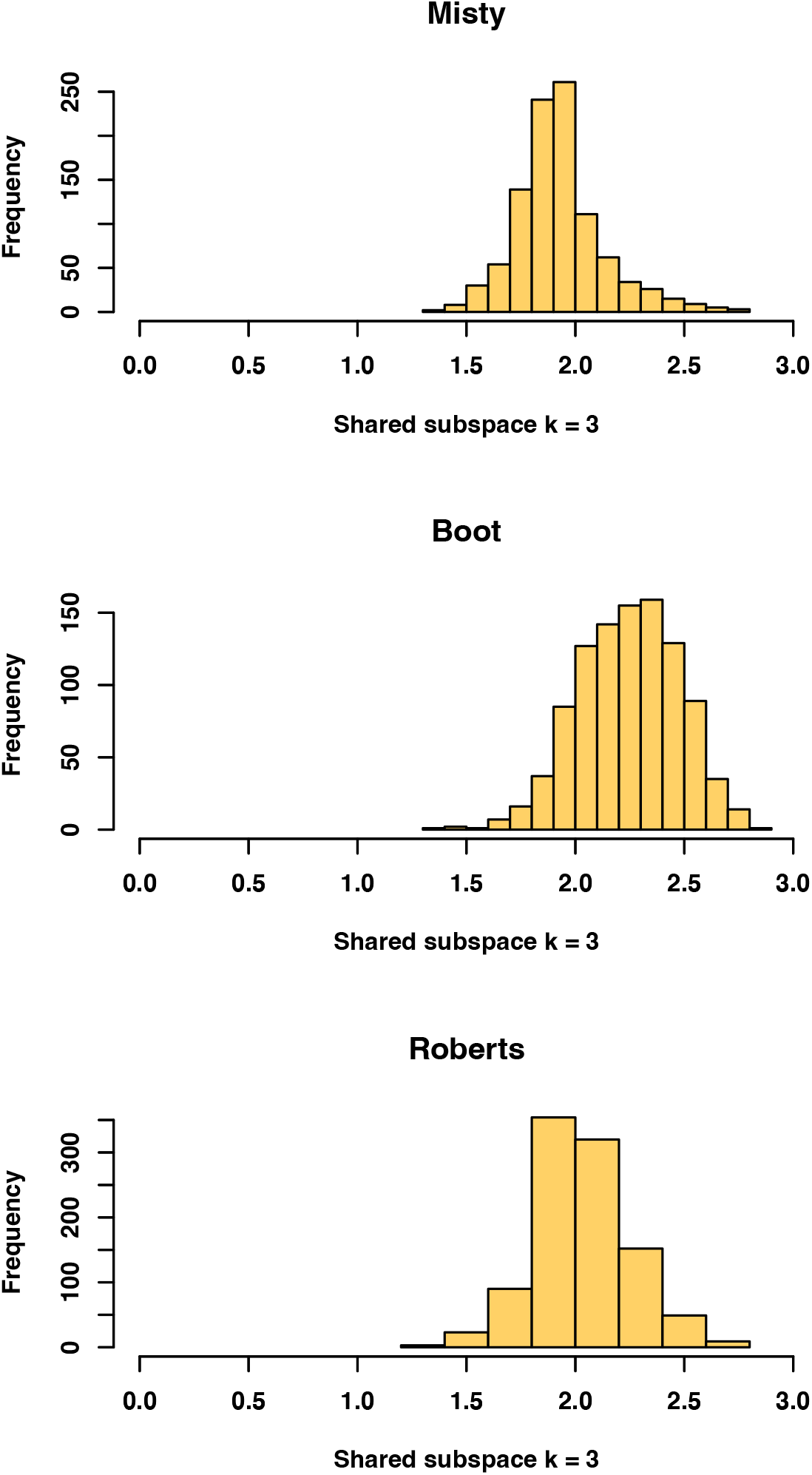
Comparing **P** matrix structure across lab-reared and wild-caught fish. Histograms show posterior distribution of Krzanowski subspace comparisons for (A), Misty (B), Boot (C), Roberts. A value of three for this statistic indicates identical matrix structure, and a value of zero indicates unrelated matrices.

